# Disease resistance of *Brassica juncea* to *Sclerotinia sclerotiorum* is established through the induction of indole glucosinolate biosynthesis

**DOI:** 10.1101/2024.01.29.577696

**Authors:** Jinze Zhang, Xu Yang, Yingfen Jiang, Hairun Jin, Kunjiang Yu, Lijing Xiao, Qingjing Ouyang, Entang Tian

## Abstract

Sclerotinia stem rot (SSR), caused by *Sclerotinia sclerotiorum*, is the main disease threat to oilseeds in Brassiceae, causing significant yield losses and reduction in oil content and quality. The studies on *S. sclerotiorum* require a great focus and extensive research on *B. juncea* compared to those on *B. napus* and *B. oleracea*. Transcriptome analysis revealed a large number of defense-related genes and response processes in *B. napus* and *B. oleracea*. However, similarities and differences in the defense responses to *S. sclerotiorum* on *B. juncea* are rarely reported. In the present study, we reported a *B. juncea* breeding line of H83 with high *S. sclerotiorum* resistance, which was used for transcriptome analysis compared to L36 with low resistance. A novel regulatory network was proposed to defend against *S. sclerotiorum* invasion in *B. juncea*. Upon infection of *S. sclerotiorum*, a series of auxin and MAPK signaling pathways were initiated within 12 h, and then defenses were activated to restrict the development and spread of *S. sclerotiorum* by inducing the massive synthesis of indole glucosinolates after 24 h. Twelve hub genes involved in the network were identified by the weighted gene co-expression network (WGCNA), which are involved in plant-pathogen interaction, signaling pathway genes, indole glucosinolate biosynthesis and cell wall formation. The hub genes were further validated by qRT-PCR. The research revealed a new resistant line of H83 against *S. sclerotiorum* and a different regulatory network in *B. juncea*, which would be beneficial for the future effective breeding of Sclerotinia-resistant varieties.

## Introduction

*Sclerotinia sclerotiorum* (Lib.) de Bary, is a very efficient plant pathogen and could infect more than 600 plant species above or below the soil surface worldwide [1]. This pathogen is primarily responsible for the development of Sclerotinia stem rot (SSR), which can lead to severe yield losses in rapeseed cultivation [2–6]. For each percent increase in SSR incidence, rapeseed yield could decrease by 0.5% for each percent increase in SSR incidence [4]. In addition, the oil content and seed quality are significantly reduced after infection [7]. Therefore, controlling the *S. sclerotiorum* epidemic represents a major challenge in rapeseed cultivation due to the pathogen’s broad host range and its ability to survive as sclerotia for long periods of time.

The mechanisms underlying the interaction between pathogens and plants have been extensively studied. The fungus *S. sclerotiorum* could acidify the environment, suppress antioxidant enzyme activities, and stimulate the generation of reactive oxygen species (ROS), simultaneously damaging host cell membranes through the secretion of oxalic acid (OA) [8]. Another arsenal of *S. sclerotiorum* is cell wall degrading enzymes (CWDEs) [5,6,9,10], which lead to the dissolution of the subcuticular epidermal walls, rapid cell death and development of necrotic symptoms. In addition, numerous small, secreted proteins of *S. sclerotiorum* have been reported to have effector-like features or effector-like activities, which are released either extracellularly or in the cytoplasm of plant cells to override host defense mechanisms and enhance fungal infections [11].

After invasion by *S. sclerotiorum*, the host defense mechanism could be activated after an attack. In *B. napus*, induction of defense related genes is often associated with the onset of resistance signaling post-pathogen infection [10,12,13]. The widespread use of RNA-sequencing (RNA-Seq) techniques has helped identify many signaling pathways, including gibberellic acid (GA), ethylene (ET), jasmonate (JA), salicylic acid (SA), auxin, and mitogen-activated protein kinase (MAPK) [13–16]. Most of these genes of signaling pathways were up regulated as response to post-pathogen infection of *S. sclerotiorum*. After a series of signal transductions, active plant defenses are induced to limit the development of pathogens. This can be achieved through by producing antimicrobial compounds and strengthening the damaged cell walls. Knocking down *BnF5H* [17] and overexpression of *cinnamoyl-coA reductase 2* [18], which regulates lignin biosynthesis, could significantly increase resistance to *S. sclerotiorum* in *B. napus*. Furthermore, monolignol biosynthesis could strengthen plant cell walls and is associated with increasing the resistance to *S. sclerotiorum* in Camelina sativa [19]. In addition, as important antimicrobial compounds in Brassiceae, the glucosinolates and their hydrolyzed products may play an important role in controlling the invasion of *S. sclerotiorum* [6,20–25]. Important antimicrobial compounds have been reported to function in *B. oleracea* [26], *B. napus* [27], and Arabidopsis [28], the increase of which could increase host resistance against *S. sclerotiorum*. Although only scant information has been reported on the function of glucosinolate in *B. juncea*.

To date, there are no immunological or highly resistant germplasms against *S. sclerotiorum*, so there is an urgent needed to screen additional genetic resources and investigate genetic mechanisms. While most work focuses on *B. napus* and *B. oleracea*, further work is needed on *B. juncea* as the fourth oilseed crop and close relative of the third oilseed crop of *B. napus*. Two methods for large-scale germplasm screening are reported, namely leaves and stems, and a close correlation between these two methods has been found [29]. Then leaf identification is a better choice for the large-scale screening of *B. juncea* germplasms, because the identification can start very early, and we have many times available for inoculation compared to the stem method. The identified germplasms could then be further confirmed by stem method. In this study, a *B. juncea* line of H83 with high resistance to *S. sclerotiorum* was reported and used to analyze its mechanism against *S. sclerotiorum* at 0 h, 12 h, 24 h, and 36 h after inoculation compred with low resistance line from L36. The work could provide us with a new case study on the mechanism of resistance to *S. sclerotiorum* in *B. juncea* except *B. napus* and *B. oleracea* in Brassiceae, which could be useful for further germplasm screening and effective breeding of Sclerotinia-resistant varieties.

## Materials and methods

### Plant materials, pathogen isolation and pathogen inoculation

The *S. sclerotiorum* strains were sampled at 18 sites in two of the nine cities in Guizhou Province, China. All *S. sclerotiorum* stains collected were isolated and confirmed to be the same by a pair of specific molecular marker (Table S1). Subsequently, the strain *gz1* was used to inoculate the *B. juncea* breeding lines to screen lines with better resistance. The isolated *S. sclerotiorum* strain *gz1* was cultured on potato dextrose agar (PDA) medium (300 g/L diced potatoes, 20 g/L sucrose and 15 g/L agar in 1 L ddH_2_O) at 25°C for activation 4 days. Then, three 3-month-old fresh young leaves from three plants of each candidate *B. juncea* breeding line were brought to the laboratory for inoculation, on which the small mycelial plugs (5 mm in diameter) cut from plates with growing fungal cultures were collected and placed. The inoculated leaves were incubated in a sealed and humidified dish at room temperature. Leaves with three biological replicates of each *B. juncea* line were sampled at 0 h, 12 h, 24 h, and 36 h after inoculation (HAI). A total of 24 samples were immediately placed in a -80°C refrigerator for RNA extraction.

### RNA extraction and sequencing

Total RNA (2μg) was extracted from the 24 treated leaves of H83 and L36 using the TRIzol kit (Invitrogen, Carlsbad, CA) according to the manufacturer’s instructions. The RNA purity of each sample was checked using a Kaiao K5500® Spectrophotometer (Kaiao, Beijing, China), and RNA integrity and concentration were measured using the RNA Nano 6000 Assay Kit for the Bioanalyzer 2100 System (Agilent Technologies, CA, USA). Subsequently, all these 24 RNA samples were sent to BioMarker Technologies (http://www.biomarker.com.cn/about-us) for cDNA library construction and Illumina deep sequencing (Wang et al. 2016).

### Data analysis and differential expression gene identification

The raw RNA sequencing data were filtered by a Perl script following the steps of the published work (Wu et al. 2016) and our previous study (Khattak et al. 2019, Wan et al. 2023). DESeq2 v1.6.3 was used for differential gene expression analysis between different samples with three biological replicates under the theoretical basis obeys the hypothesis of the negative binomial distribution for the count value. The p value was corrected using the BH method. Genes with p≤0.05 and |log2_ratio|≥1 were identified as differentially expressed genes (DEGs) [30]. The obtained DEGs were further annotated using Gene Ontology (GO, http://geneontology.org/) and analyzed by KEGG (Kyoto Encyclopedia of Genes and Genomes, http://www.kegg.jp/) [31,32]. GO enrichment of DEGs was implemented by the hypergeometric test, where the p value is calculated and adjusted to produce the q-value, and the data background is the genes in the whole genome. GO terms with q<0.05 were considered significantly enriched. GO enrichment analysis was used to determine the biological functions of the DEGs. The KEGG enrichment of the DEGs was determined by the hypergeometric test, in which the p value was adjusted by multiple comparisons to obtain the p value. KEGG terms with q<0.05 were considered significantly enriched.

### Gene co-expression network analysis and visualization

To better identify the core pathway and hub genes, all genes from RNA-seq were analyzed to construct the gene co-expression networks using the R package WGCNA [33]. The genes with kME >0.7 were selected as member of each module, which were all divided into 14 modules by dynamic tree cutting and merged dynamic methods using *β*=22 as the soft threshold power. Module-trait associations were estimated using Pearson’s correlation coefficients, and the correlation is reflected in the color depth in the heatmap. All genes in the key modules were used to build a protein-protein interaction network (PPI) via STRING (v12.0) [34], and hub genes were consistently identified via four algorithms (degree, MNC, stress, and MCC) of cytoHubba in Cytoscape_v3.9.1 [35]. Each node represents a gene connected to a different number of genes. The gene that is associated with a larger number of genes is characterized by a larger size and is more important for its interaction with a large number of genes.

### Quantitative real-time PCR (qRT-PCR) analysis

Quantitative real-time PCR (qRT-PCR) was used to confirm the RNA-seq results and analyze the expression level of the identified hub genes. A total of 2 ug of the total RNA from each sample were used to synthesize the cDNA using PrimeScript RT Reagent Kit with gDNA Eraser (Takara, Dalian, China) according to the manufacturer’s instructions. Twelve gene-specific primers for qRT-PCR were designed based on sequencing sequences of the identified hub genes using Primer Premier 6.0. Real-time PCR was performed using SsoAdvanced^TM^ Universal SYBRR Green Supermix (Hercules, CA) according to our previous research [36,37]. The 2^-ΔΔCt^ algorithm was used to calculate the relative level of gene expression. The *β-actin* gene was used as an internal control. All qRT-PCR was performed with three biological replicates and run on a Bio-Rad CFX96 Real Time System (Bio-Rad, Hercules, CA, USA). The primers used in these experiments are listed in (Table S1).

## Results

### Isolation, purification, and inoculation of *S. sclerotiorum*

To better study the species of *S. sclerotiorum* in Guizhou Province, China and isolate the pathogen for resistant rapeseed breeding. We collected a total of 18 samples from the night city area, i.e. at least two samples for each city (Fig. 1A). The samples were isolated in the laboratory and cultured on the PDA culture medium (Fig. 1B). DNA from all 18 samples were extracted and amplified with a primer pair (Fig. 1C) normally used to identify the *S. sclerotiorum* type. The amplified bands (Fig. 1D) were purified and sent for sequencing. With the exception of a sample called *gz12*, which detected an A to G mutation (Fig. 1E), all other sequences were completely consistent. After searching the NCBI database, all these pathogen samples were determined to be *S. sclerotiorum*. The pathogen sample from *gz1* was used for subsequent inoculation and analysis.

**Figure 1.**
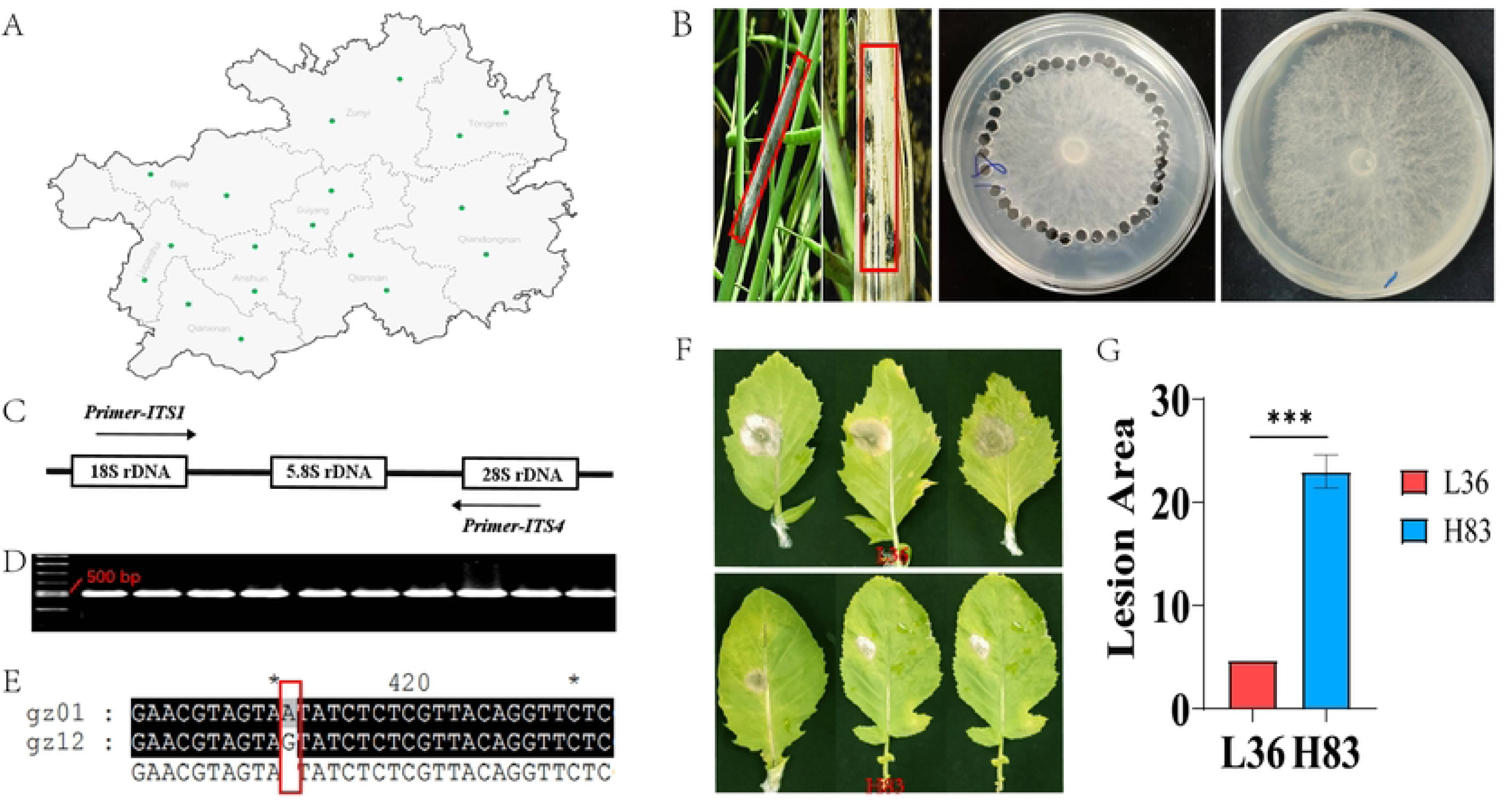
Isolation, identification and inoculation of *S. sclerotiorum*. The distribution of the sites for sampled *S. sclerotiorum* stains (A), the isolation of the sampled stains (B), the primer position (C), amplifying results (D) and a A to G mutation in strain *gz12* (E), the inoculation of *B. juncea* lines of H83 and L36 (F), the analysis of lesion area between H83 and L36.

To examine the lines with better resistance to *S. sclerotiorum* in *B. juncea*, *gz1* was used to inoculate the leaves of 200 *B. juncea* lines (data not shown). Two samples named L36 with low resistance and H83 with high resistance to *S. sclerotiorum* (Fig. 1F) were selected for transcriptome analysis. The average lesion area of L36 is 23.00 cm^2^, which is significantly larger than 4.71 cm^2^ of H83 (p=000) (Fig. 1G).

### Identification and characterization of differential expression genes

Three developing leaves for each line of L36 and H83 were inoculated with the isolated *S. sclerotiorum* pathogen strain *gz1*, and sampled for transcriptome analysis at 0 h, 6 h, 12 h, and 24 h. A total of 503,358,531 clean reads, 150,748,528,997 clean bases and 150.72 Gb clean data were obtained from the 24 leaf samples through the RNA-seq analysis (Table S2). Three replicates of each phase and line were clustered together (Fig. 2A). The percentage of the Q30 base ranged from 93.09% to 94.35%, and the average GC content was 46.30% (Table S2). The clean reads of each sample were compared with the designated *B. juncea* reference genome (v 1.5) [38], and the comparison efficiency ranged from 92.09% to 93.43% (Table S3).

**Figure 2.**
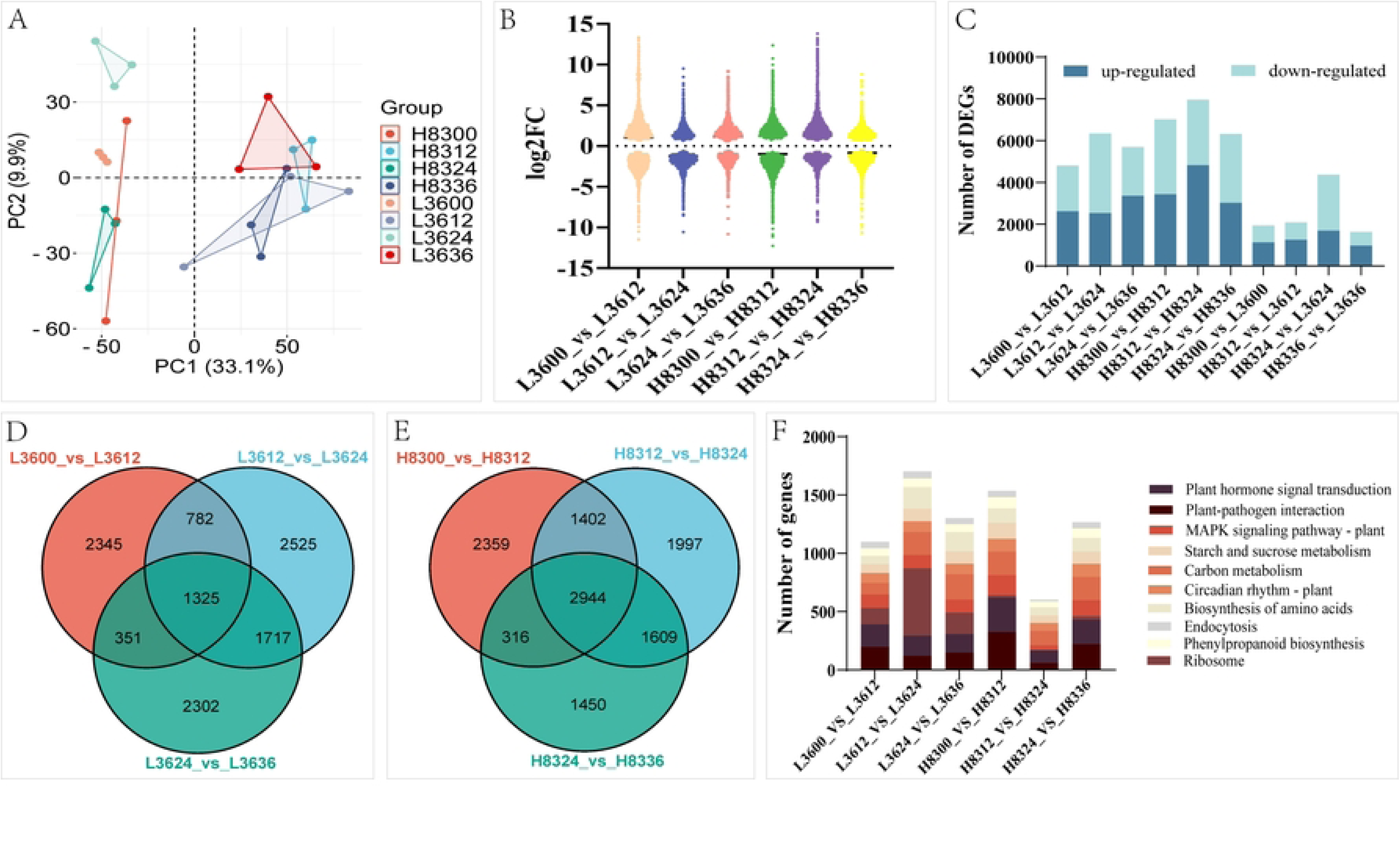
Analysis of the RNA-seq results. The PCA analysis for the 24 sequenced samples (A), the statistics for the log2FoldChange of different comparison groups, the number of DEGs in different comparison groups, the Venn diagram indicating the common DEGs among different comparison groups of L36 (D) and H83 (E), the number of DEGs in different KEGG pathways within different comparison groups.

The expression level of different phases and lines were compared with |log2 (FoldChange)| > 1 and p < 0.05 as the standard. The fold values varied between different comparison groups (Fig. 2B). For line L36, 4,803 DEGs (2,617 upregulated and 2,186 downregulated) between 0 h and 12 h after inoculation, 6,349 DEGs (2,541 upregulated and 3,808 downregulated) between 12 h and 24 h after inoculation, and 5,695 DEGs (3,365 upregulated and 2,330 downregulated) between 24 h and 36 h after inoculation, were detected (Fig. 2C; Table S4). Among these three comparison groups of L36, 1,325 common DEGs were detected (Fig. 2D). For line H83, 7,021 DEGs (3,440 upregulated and 3,581 downregulated) between 0 h and 12 h after inoculation, 7,952 DEGs (4,828 upregulated and 3,124 downregulated) between 12 h and 24 h after inoculation, and 6,319 DEGs (3,025 upregulated and 3,294 downregulated) between 24 h and 36 h after inoculation, were detected (Fig. 2C; Table S4). Among these three comparison groups of H83, 1,325 common DEGs were detected (Fig. 2D). There are still 832 common DEGs among these six comparison groups.

The KEGG pathways of these DEGs in different comparison groups were also analyzed. The first three pathways of these were plant hormone signal transduction for 1,142 genes, plant-pathogen interaction for 1,069 genes, and carbon metabolism for 1,040 genes (Fig. 2F; Table S5).

### Weighted correlation network analysis for key modules and pathways

To identify *S. sclerotiorum*-induced hub genes underlying pathogen resistance regulatory pathways, we constructed a weighted gene co-expression network (WGCNA) based on RNA-seq data from L36 and H83 leaf samples infected with *S. sclerotiorum* using the R package WGCNA [33]. After filtering out the low-abundance and low-variability genes, a total of 9,679 genes were screened out (Table S6) and imported into the WGCNA software package for the analysis. Through WGCNA, a co-expression network was constructed and in which 14 modules were identified via the dynamic tree cutting and merged dynamic methods using *β*=22 as the soft threshold power (Fig. 3A-3C). The genes with kME>0.7 were chosen as the member of each module, and the number of which is shown in Fig. 3D. The divided module shows tight correlations with the traits, since the samples inoculated at the same time were clustered together (Fig. 3E), and not determined by genotypes. Eigengene–trait correlation analysis revealed that six modules of MEbisque4, MElightblue3, MEsalmon4, MEdarkred, MEpalevioletred1, and MElightpink3 exhibited a positive contribution to *S. sclerotiorum* resistance in H83 (Fig. 3F). Therefore, these six modules were used for subsequent analysis.

**Figure 3.**
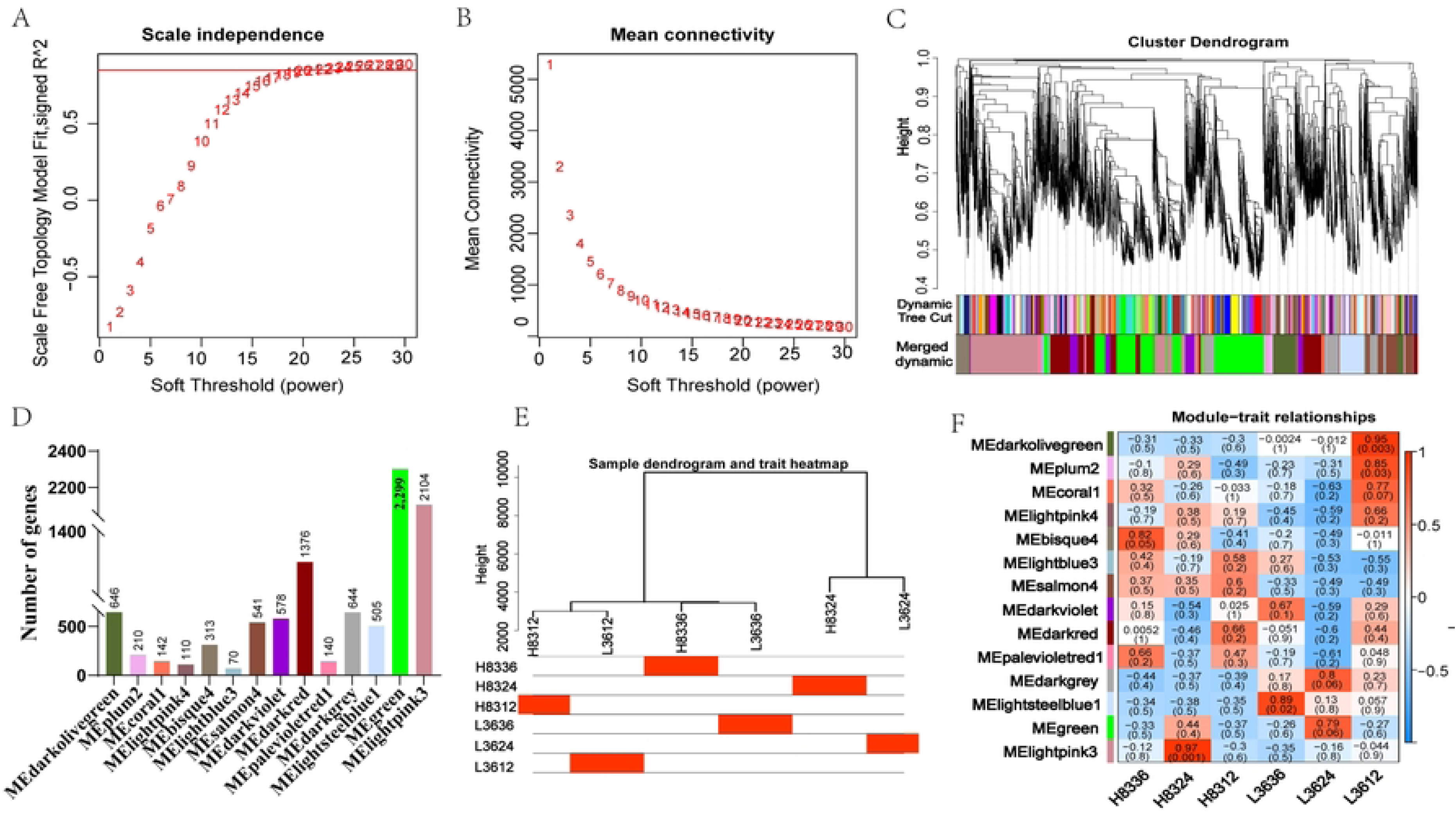
Results of WGCNA analysis. The soft threshold power of β=22 (A and B), the dynamic tree cutting and merged dynamic modules (C), the number of genes in each module (D), the sample dendrogram and trait heatmap of the 6 samples (E), the module-trait relationships of the divided 14 modules (F). The redder color indicates the high correlation between traits and module, the bluer color indicates the more negative correlation (F).

KEGG enrichment analysis was performed to biologically characterize the key pathways in the selected six modules (Fig. 4). MEbisque4 is one of the key modules and showed a significant positive correlation (r=0.82, p=0.05) with the *S. sclerotiorum* resistance in H86 at 36 h, in which glucosinolate biosynthesis, glyoxylate and dicarboxylate metabolism, and carbon fixation in photosynthetic organisms were the top three pathways (Fig. 4A). For the MElightblue3 module, phenylpropanoid biosynthesis, plant-pathogen interaction, and spliceosome were the top three pathways (Fig. 4B). For the MEsalmon4 module, alpha-linolenic acid metabolism, nucleotide excision repair, and mismatch repair were the top three pathways (Fig. 4C). For the MEdarkred module, circadian rhythm-plant, plant hormone signal transduction, and sulfur metabolism were the top three pathways (Fig. 4D). For MEpalevioletred1 module, fatty acid elongation, arachidonic acid metabolism, and glycine, serine and threonine metabolism were the top three pathways (Fig. 4E). Furthermore, MElightpink3 is another key module and showed a significant positive correlation (r=0.97, p=0.0001) with *S. sclerotiorum* resistance in H86 at 24 h, in which plant-pathogen interaction, glutathione metabolism, and MAPK signaling pathway-plant were the top three pathways (Fig. 4F).

**Figure 4.**
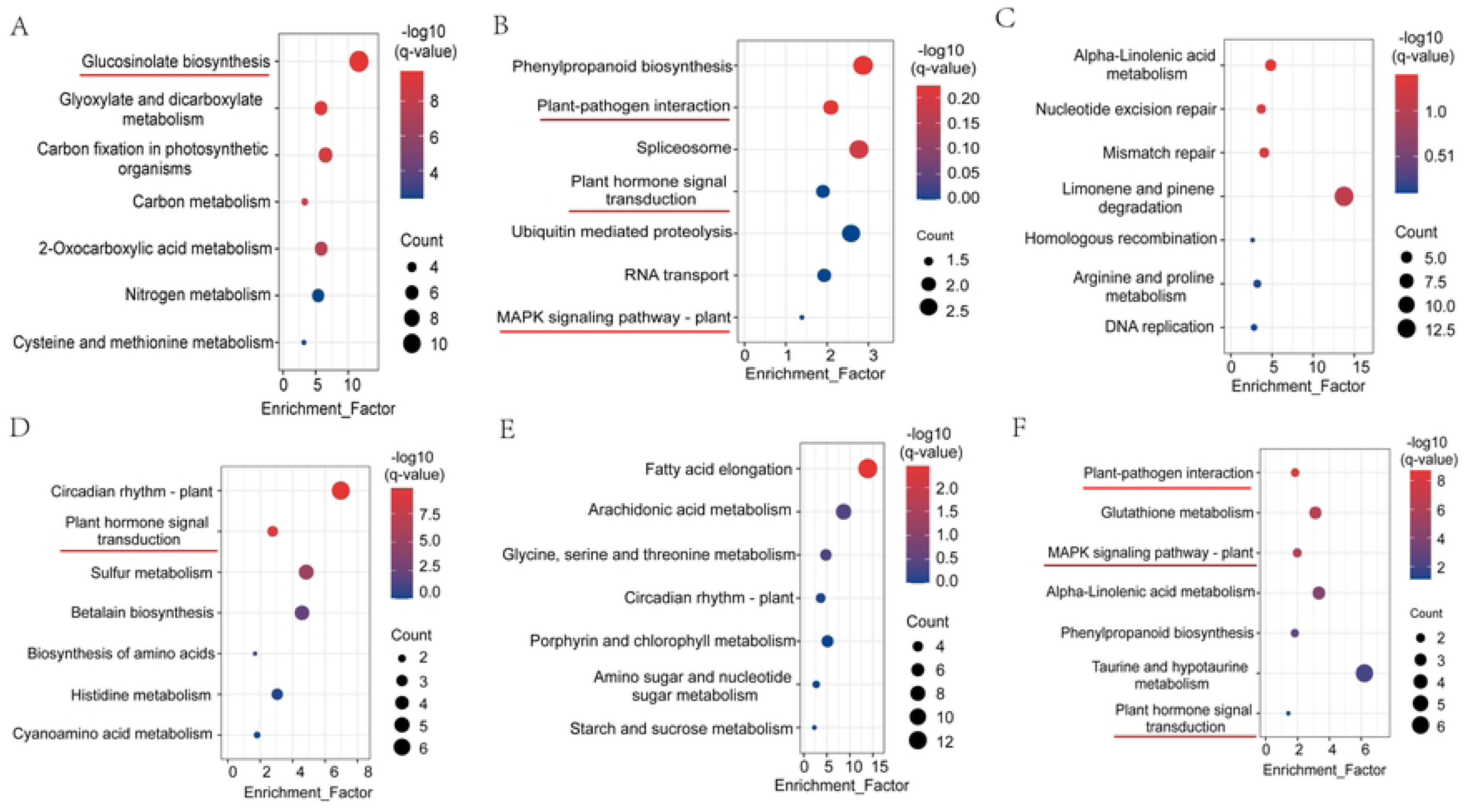
The KEGG pathways of the six selected modules. The bubble chart of selected modules for MEbisque4 (A), MElightblue3 (B), MEsalmon4 (C), MEdarkred (D), MEpalevioletred1 (E), and MElightpink3 (F). The bigger circle indicates higher enrichment, and the redder color indicates the higher value of -log10(q-value).

### Analyzing the signaling pathway and glucosinolate metabolism pathway

According to WGCNA analysis, MEbisque4 and MElightpink3 are the two main modules, in which MAPK signaling pathway, plant hormone signal transduction, and glucosinolate biosynthesis play an important role in resistance to *S. sclerotiorum*. For plant hormone signal transduction, the number of DEGs were compared in different comparison groups (Fig. 5A), involving the signaling molecules of slicylic acid, jasmonic acid, gibberellin, ethylene, cytokinine, brassinosteroid, auxin and abscisic acid. In line L36, more auxin DEGs were induced at 12 h that at 24 h and 36 h. While the highest number of DEGs for auxin was induced at 12 h in L83 significantly higher than that at 12 h in L36. For the MAPK signaling pathway, the number of induced DEGs was significantly higher (p=0.000) in H83 than in L36 at 12 h, 24 h and 36 h (Fig. 5B), indicating the important role of this pathway in its resistance to *S. sclerotiorum*.

**Figure 5.**
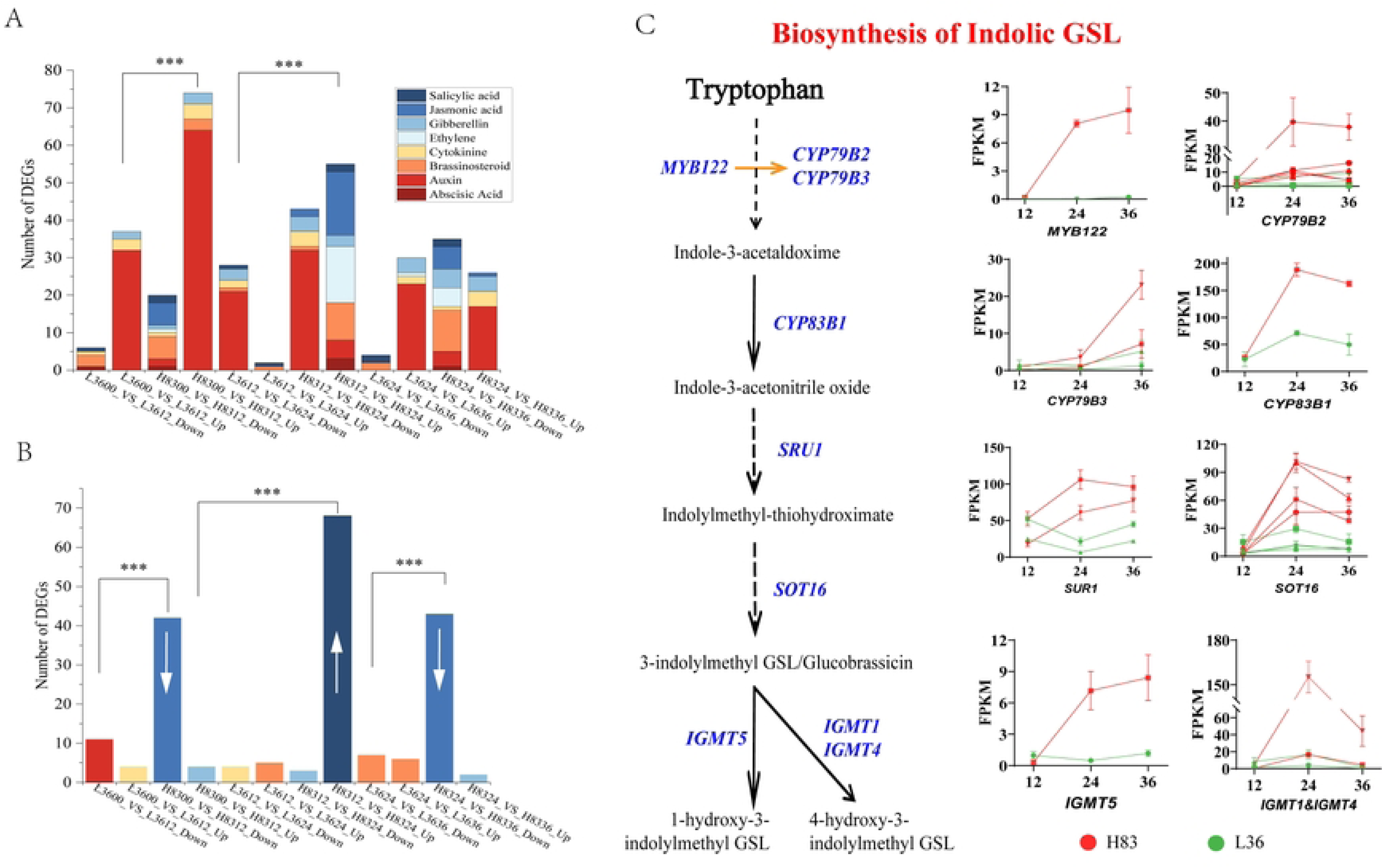
Statistics of the number of DEGs involved in key signaling pathways and the gene expression level involved in the indole glucosinolate biosynthesis. The number of DEGs for hormone signaling pathway (A) and MAPK signaling pathway (B). The expression level for 9 genes involved in indole glucosinolate biosynthesis (C).

The KEGG pathway of glucosinolate (GSL) biosynthesis was mainly related to the biosynthesis of indolic GSLs (Fig. 5C). From tryptophan to 1-hydroxy-3-indolylmethyl GSL and 4-hydroxy-3-indolylmethyl GSL, the expression level of orthologous genes of *MYB122* (*BjuB030170*), *CYP79B2*, (*BjuO008644*, *NewGene_9671*, *BjuA002613*, *BjuB038533* and *NewGene_15815*), *CYP79B3* (*BjuA015924* and *BjuA015925*), *CYP83B1* (*BjuA028230*), *SRU1* (*BjuB029587* and *BjuB029587*), *SOT16* (*BjuA043506*, *BjuB030167*, *BjuB010311* and *BjuA005541*), *IGMT1* (*NewGene_13593*), *IGMT4* (*BjuB043479*) and *IGMT5* (*NewGene_5430*) in H83 increased significantly compared to L36 at the same time (Fig. 5C). The results show that the increase started from 24 h and continued until 36 h after inoculation of *S. sclerotiorum*.

### Identification of hub genes from the key modules

All genes in the key modules MEbisque4 and MElightpink3 were used to build a protein-protein interaction (PPI) network via STRING (v12.0) [34], and hub genes were consistently identified via four algorithms (degree, MNC, stress, and MCC) of cytoHubba in Cytoscape_v3.9.1 [35]. The top six gene scores with functional annotations for each of the two key modules of MEbisque4 and MElightpink3 were considered as hub genes (Table 1). To further localize the hub genes, all DEGs associated with these 12 hub genes within these two modules were used to construct a PPI network (Fig. 6A and 6B). Apparently, a high correlation was observed in the network between the hub genes in each module, suggesting that they are synergistically involved in regulating resistance to *S. sclerotiorum*. The important essential roles of these hub genes in these two networks were represented by the larger circle and redder color (Figs. 6A and 6B).

**Figure 6.**
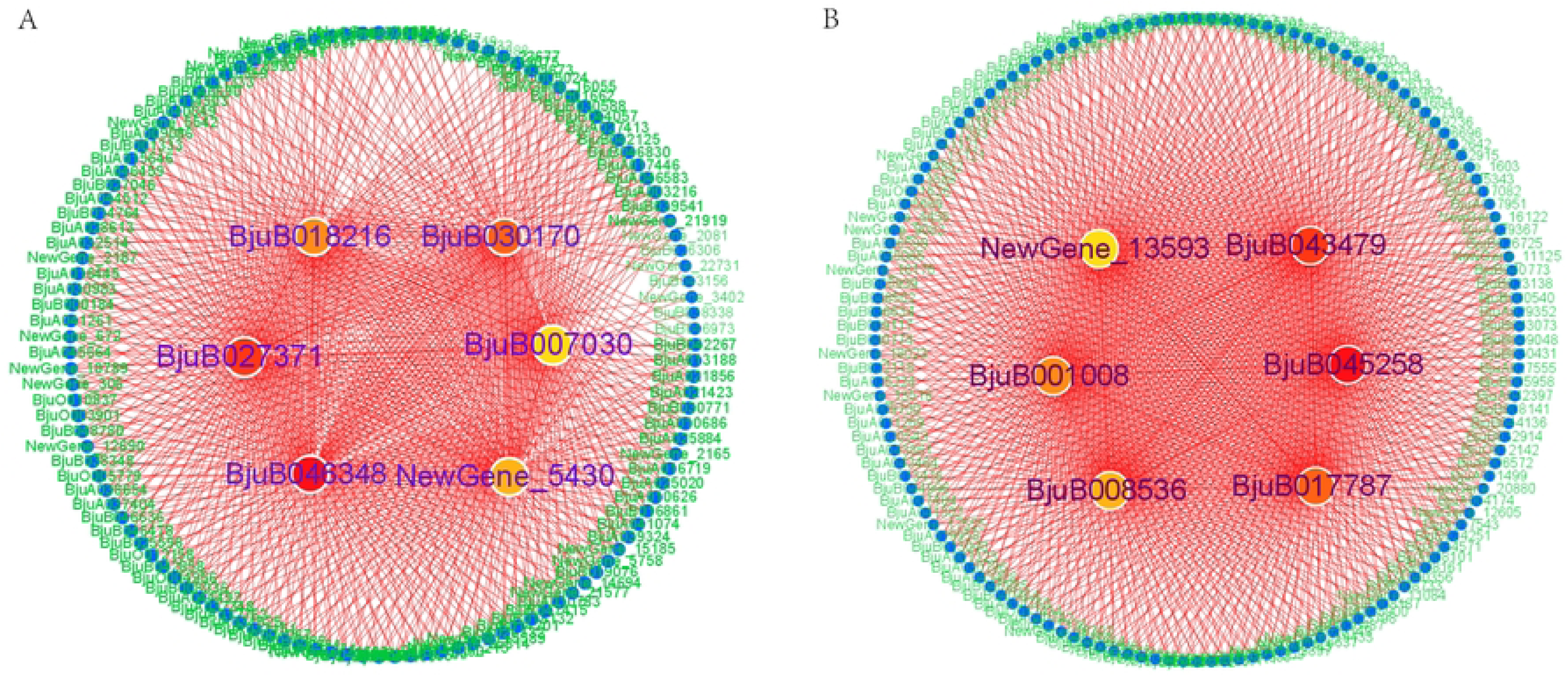
Gene co-expression network analysis and visualization. Screening of hub genes in the key modules of MEbisque4 (A) and MElightpink3 (B) was done via four algorithms (degree, MNC, stress, and MCC) from cytoHubba in Cytoscape_v3.9.1. The identified 12 hub genes and their interaction with other genes were shown, the number of lines indicates their stronger correlation with other genes. The larger circle indicates high correlation with other genes, and the redder color indicates higher importance of the gene.

**Table 1.**
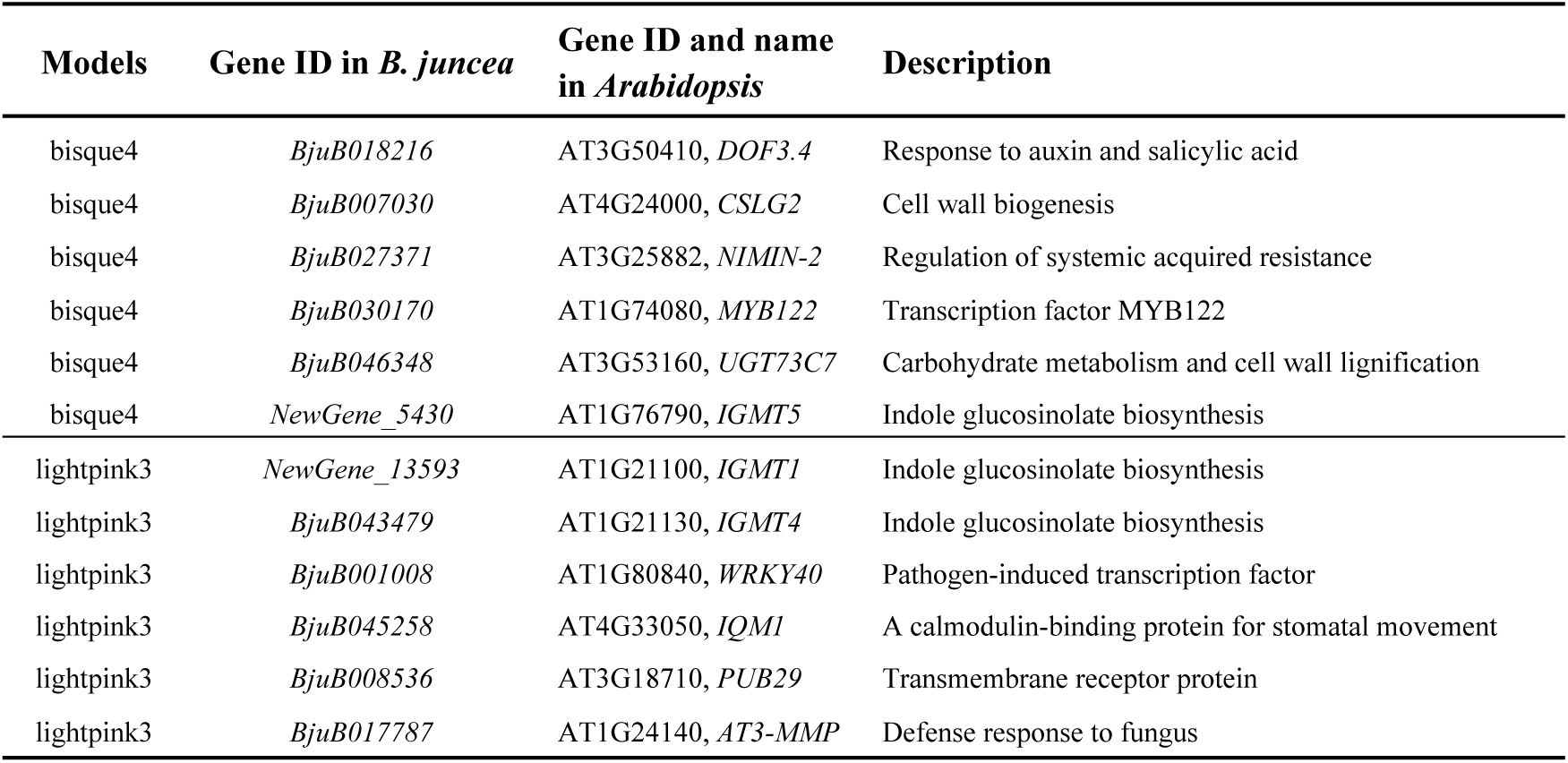
The Information of hub genes in two key modules.

Among the 12 hub genes within the MEbisque4 and MElightpink3 modules were further functionally identified. Six genes of *BjuB018216* (*DOF3.4*), *BjuB027371* (*NIMIN-2*), *BjuB001008* (*WRKY40*), *BjuB045258* (*IQM1*), *BjuB008536* (*PUB29*) and *BjuB017787* (*AT3*-*MMP*) for plant-pathogen interaction and signaling pathway were identified. Four genes of *BjuB030170* (*MYB122*), *NewGene_5430* (*IGMT5*), *NewGene_13593* (*IGMT1*), and *BjuB043479* (*IGMT4*) were identified for indole glucosinolate biosynthesis. In addition, another two additional genes of *BjuB007030* (*CSLG2*) and *BjuB046348* (*UGT73C7*) were identified, which are responsible for cell wall formation.

### qRT-PCR validation of the hub genes

The expression level of these 12 hub genes were further validated by qRT-PCR (Fig. 7). The expression level for the hub genes of *BjuB018216* (*DOF3.4*) (Fig. 7A), *BjuB030170* (*MYB122*) (Fig. 7D), and *NewGene_5430* (*IGMT5*) (Fig. 7F) in L36 is significantly higher at 12 h and significantly lower at 24 h and 36 h compared to H83. The expression level for the hub genes of *BjuB007030* (*CSLG2*) (Fig. 7B), *BjuB027371* (*NIMIN-2*) (Fig. 7C), *BjuB046348* (*UGT73C7*) (Fig. 7E), and *BjuB043479* (*IGMT4*) (Fig. 7H) showed no significant difference at 12 h and significantly higher at 24 h and 36 h in H83 compared to L36. The expression level of genes of *NewGene_13593* (*IGMT1*) (Fig. 7G), *BjuB001008* (*WRKY40*) (Fig. 7I), *BjuB045258* (*IQM1*) (Fig. 7J), *BjuB008536* (*PUB29*) (Fig. 7K) and *BjuB017787* (*AT3*-*MMP*) (Fig. 7L) in H83 at 24 h were significantly higher than that in L36, and there was no significant difference in other phases between H83 and L36.

**Figure 7.**
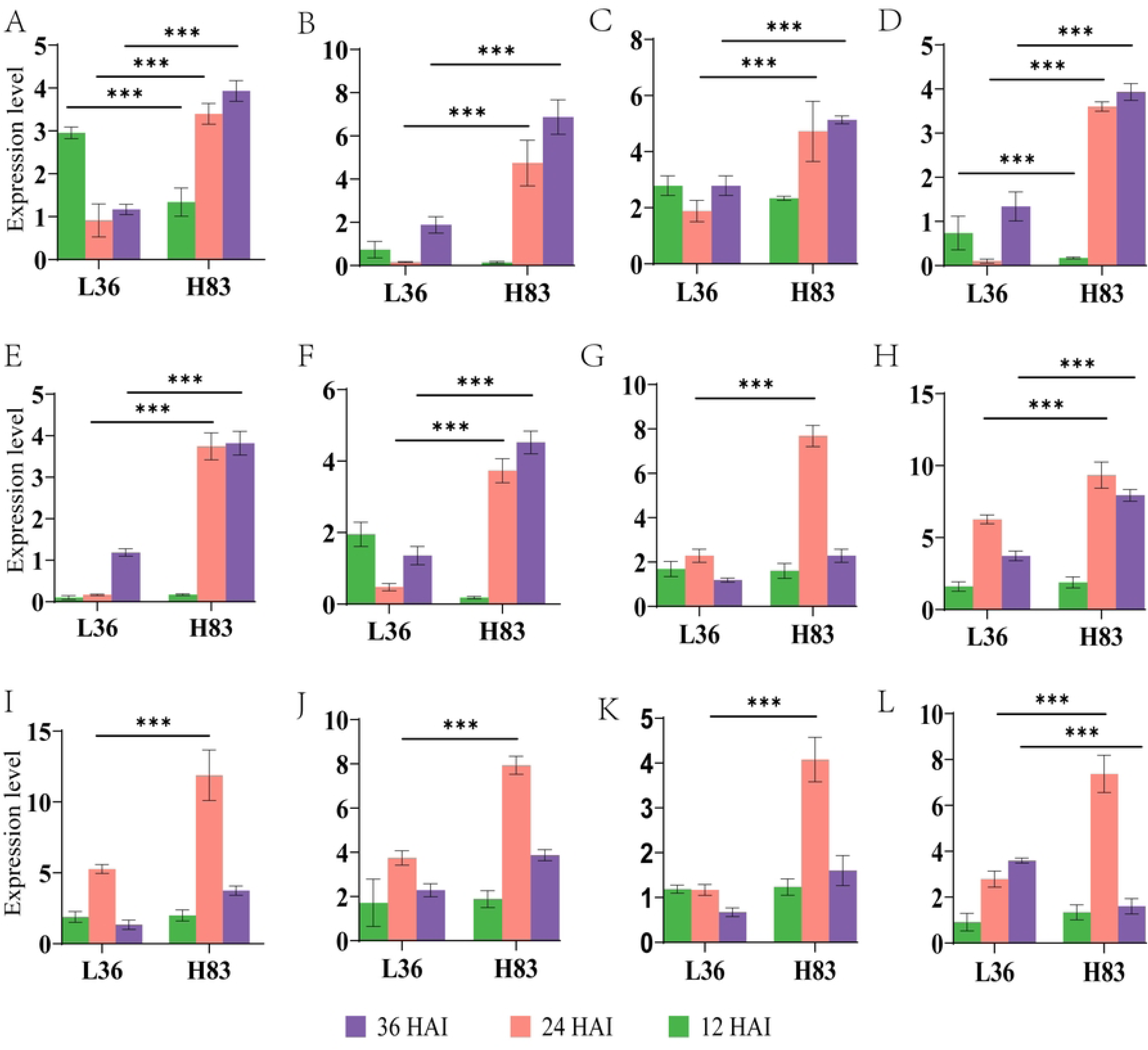
Validation of the 12 hub genes via real-time qRT-PCR. The expression level of BjuB018216 (Fig. 7A), (B)BjuB007030 (Fig. 7B), (C)BjuB027371 (Fig. 7C), (D)BjuB030170 (Fig. 7D), (E) BjuB046348 (Fig. 7E), (F)NewGene_5430 (Fig. 7F), (G)NewGene_13593 (Fig. 7G), (H)BjuB043479 (Fig. 7H), (I) BjuB001008 (Fig. 7I), (J)BjuB045258 (Fig. 7J), (K)BjuB008536 (Fig. 7K), and (L)BjuB017787 (Fig. 7L) was shown, blue, red and green color indicate the 12, 24 and 36 hours after innovation (HAI). * indicates the significant difference at p<0.05, ** indicates the significant difference at p<0.001, ** indicates the significant difference at p=0.000.

## Discussion

*S. sclerotiorum* is one of the world’s most important fungal diseases, which can lead to enormous yield losses in many species, especially oil plants in Brassiceae. Unfortunately, no immune or highly resistant germplasms have been found so far. This means that screening more genetic resources and studying genetic mechanisms were urgently needed to develop an effective strategy for *Sclerotinia* resistance breeding. In *B. napus*, much work has been done to screen resistant germplasm, although the quality is limited. However, as an import relative of *B. napus*, some lines with relatively higher resistance to *S. sclerotiorum* were identified in *B. oleracea* and increased the resistance of *B. napus* via distant hybridization [39,40]. Although *B. juncea* is the fourth largest oilseed crop in the world, there is particularly little research on it. In the present study, we reported a *B. juncea* breeding line of H83 with relatively high levels of *S. sclerotiorum* resistance compared to L36 and analyzed its transcriptomic changes after 12, 24, and 36 h inoculation. Our results revealed the specific defense responses in *B. juncea* and a complex and coordinated gene regulatory network that confers resistance differences between genotypes, thereby expanding our knowledge on how to increase *S. sclerotiorum* resistance in the future.

Facing the fungal invasion, the expression level of genes in the signaling pathway usually increases rapidly. While the main signaling pathway usually varies with the genotype difference. In *B. napus*, the genes involved in the regulating GA and ET synthesis are significantly up regulated in the resistant line [14]. In another report, JA and ET were significantly up regulated signaling pathway in *B. napus* [13]. It was once reported that JA pathway was repressed at 24 h after inoculation, while SA was activated in *B. napus* [15]. In the present study, almost all types of genes involved in the above-mentioned signaling pathway were activated in L36 and H83. While the main functional signaling pathways were auxin and MAPK, in which the number of DEGs was highest and showed a significant increase in H83 compared to L36 from 12 h after inoculation. Therefore, the functional signaling pathway in *B. juncea* might be a new signaling pathway compared to *B. napus*. As an important hub gene reported in this study, *BjuB018216* (*DOF3.4*) may play an important role in the defense against *S. sclerotiorum* in *B. juncea* as a gene response to auxin and salicylic acid.

Glucosinolates are natural plant products that play a role in the defense against herbivores and pathogens and whose biosynthesis could be significantly induced by defense signaling pathways [41]. In one study, aliphatic glucosinolate biosynthesis could be significantly induced by glucose signaling molecules [42], which are normally synthesized largely in the face of invasion by herbivores and pathogens. The indole glucosinolates could also be significantly induced by 3- to 4-fold by exogenous MeJA treatment, and the corresponding Trp-metabolizing genes *CYP79B2* and *CYP79B3* were both strongly induced [41]. In particular, the author found that the indole glucosinolate N-methoxy-indol-3-ylmethylglucosinolate accumulated 10-fold in response to MeJA treatment, whereas 4-methoxy-indol-3-ylmethylglucosinolate accumulated 1.5-fold in response to 2,6-dichloro-isonicotinic acid. In the present study, the expression level of 9 genes including *CYP79B2* and *CYP79B3* for indole glucosinolate biosynthesis was found to be significantly increased, suggesting that the defense signaling pathway is significantly induced. In addition, four indole glucosinolate biosynthesis genes of *BjuB030170* (*MYB122*), *NewGene_5430* (*IGMT5*), *NewGene_13593* (*IGMT1*), and *BjuB043479* (*IGMT4*) were identified as hub genes, further supporting their important roles in the defense of *S. sclerotiorum* in *B. juncea*.

Based on these results, we predict a potential network for defense against *S. sclerotiorum* invasion in *B. juncea* (Fig. 8). Upon infection of *S. sclerotiorum*, a series of auxin and MAPK signal pathways were initiated, and its defenses were activated to limit the development and spread of *S. sclerotiorum* by inducing the massive synthesis of indole glucosinolates.

**Figure 8.**
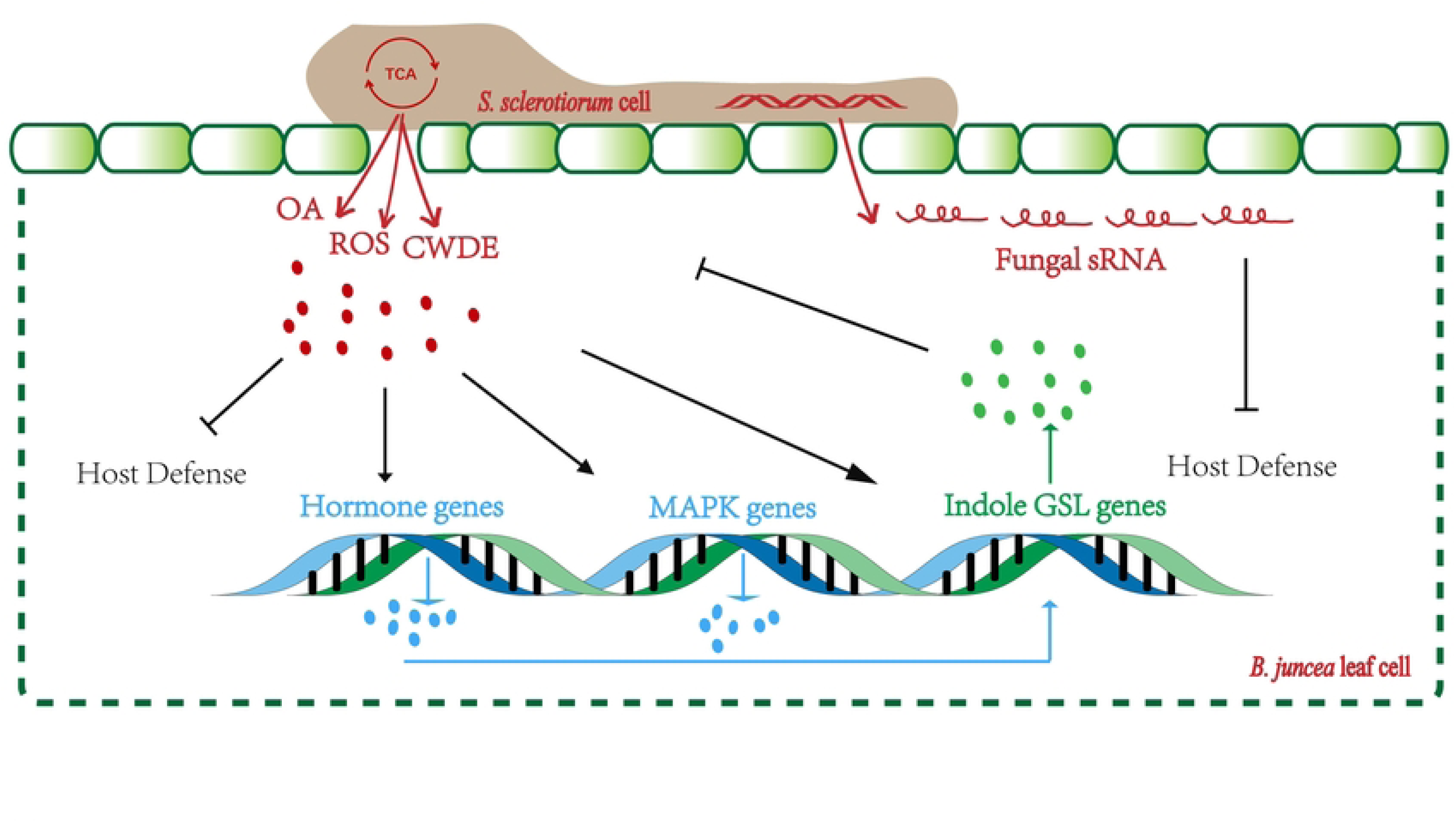
A predicted potential network for defense against *S. sclerotiorum* invasion in *B. juncea*. The three arsenals of *S. sclerotiorum* in damaging the host cell and restrict the host defense. Upon infection of *S. sclerotiorum*, a series of auxin and MAPK signal pathways were initiated, and its defenses were activated to limit the development and spread of *S. sclerotiorum* by inducing the massive synthesis of indole glucosinolates.

Further investigations are required to clarify the damage caused by *S. sclerotiorum*, particularly for *B. juncea*. *B. juncea* has many resistance genes in the B genome that have been overlooked by most experts and may provide a surprising answer to resistance against *S. sclerotiorum*. The H83 line with high resistance to *S. sclerotiorum* described in the present study would be used in the future to improve commercial varieties of *B. juncea* through intraspecific hybridization and *B. napus* through interspecific hybridization.

## Acknowledgments

This work was funded by the Scientific and Technological Key Program of Guizhou Province (No. Qiankehezhicheng [2022] Key 031), the National Natural Science Foundation of China (Grant No. 32160483 and 32360497), and the Post-Funded Project for National Natural Science Foundation of China from Guizhou University (No. [2023]093), Key Laboratory of Molecular Breeding for Grain and Oil Crops in Guizhou Province (Qiankehezhongyindi [2023]008), Key Laboratory of Functional Agriculture of Guizhou Provincial Higher Education Institutions (Qianjiaoji [2023] 007).

## Author contributions statement

ET designed the experiment; JZ and XY performed the experiments. JZ, XY, YJ, HJ, KY, LX, OO and ET analyzed the data, wrote and revised the manuscript. All authors read and approved the manuscript.

## Data Availability Statement

The data presented in this study are openly available in NCBI, reference number PRJNA1069335.

## Compliance with ethical standards

**Conflict of interest** The authors declare no competing financial interests.

**Ethical standards** The experiments were performed in compliance with the current laws of China.

## Supporting information

**Table S1 Primer sequence used in the present study.**

**Table S2 The basic information of the RNA-seq.**

**Table S3 Obtained reads were analyzed by alignment against *B. juncea* reference genomes.**

**Table S4 Statistics of the DEGs among different comparison groups.**

**Table S5 Number of genes involved in different KEGG pathways.**

**Table S6 Genes for WGCNA analysis.**

